# Latent factor modelling of scRNA-seq data uncovers novel pathways dysregulated in cell subsets of autoimmune disease patients

**DOI:** 10.1101/853903

**Authors:** Giovanni Palla, Enrico Ferrero

## Abstract

Latent factor modelling applied to single-cell RNA-sequencing (scRNA-seq) data is a useful approach to discover gene signatures associated with cell states. However, it is often unclear what method is best suited for specific tasks and how latent factors should be interpreted from a biological perspective.

Here, we compare four state-of-the-art methods and explore their stability, predictive power and coverage of known biology. We then propose an approach that leverages the derived latent factors to directly assign pathway activities to specific cell subsets. By applying this framework to scRNA-seq datasets from biopsies of rheumatoid arthritis and systemic lupus erythematosus patients, we discover both known and novel disease-relevant gene signatures in specific cellular subsets in a fully unsupervised way. Focusing on rheumatoid arthritis, we identify an inflammatory Oncostatin M receptor signalling signature active in a subset of synovial fibroblasts and dysregulation of the GAS6 - MERTK axis in a subset of synovial monocytes with efferocytic function.

Overall, we provide insights into strengths and weaknesses of latent factors models for the analysis of scRNA-seq data, we develop a framework to identify cell subtypes in a function- or phenotype-driven way and use it to identify novel pathways dysregulated in rheumatoid arthritis.

## Introduction

Single-cell RNA-sequencing (scRNA-seq) is a powerful technique that enables gene expression measurements in thousands of individual cells. Resolving cellular heterogeneity by scRNA-seq has enabled groundbreaking discovery in the biomedical domain, such as finding key disease drivers in cancer^1–3^, neurodegeneration^4,5^ and immune-mediated diseases^6–9^. From a data analysis standpoint, a crucial step in a standard scRNA-seq pipeline^10^ is clustering, where discrete cell populations sharing a common transcriptional profile are defined. These cell clusters are used in a variety of downstream analyses, such as differential expression^11–13^, compositional analysis^14^ and cellular interaction analysis^15–17^. Differential expression is fundamental to aid the phenotypic identification of cell types, usually performed by means of a hybrid approach that entails prior knowledge of the biological system and gene set enrichment analysis. An alternative, cluster-free approach to phenotypic identification of cellular states is trajectory analysis^18^, which aims to derive differentiation processes by using a pseudo-temporal ordering of single cells. However, in addition to identity- and differentiation-specific activities, transcriptional programs entail a variety of cellular processes such as metabolism, growth, stress and cell signalling, which are not necessarily captured by these approaches. Nevertheless, such expression programs are of great interest in a disease setting, where several communicating cell populations might act in the same dysregulated pathway. Thus, an in-depth characterization of such pathogenic signalling cascades at single-cell resolution is of great interest from a disease understanding perspective.

Latent factor models aim to decompose the global expression profile in its underlying transcriptional programs^19^. These models project both genes and cells in a low-dimensional space, with latent dimensions approximating cells’ transcriptional programs and summarizing the contributions of several genes. Standard matrix factorization approaches, such as principal component analysis (PCA), nonnegative matrix factorization (NMF) and independent component analysis (ICA), have been widely applied to scRNA-seq data^1,2,20^. Nevertheless, novel methods have been developed that account for the specificities of single-cell data, using meaningful prior distributions and enforcing sparsity^21–26^. A major challenge of these approaches is the determination of the number of latent dimensions to use. Despite a few heuristics having been proposed^20,27^, it is unclear whether these strategies could be applied effectively to datasets with different characteristics, and whether such heuristics are appropriate for different tasks. Furthermore, it has been shown that different biological processes are captured at different dimensionalities of the latent space^28^, suggesting that approaches considering a varying number of latent dimensions could be more robust in recapitulating the underlying biological hierarchies of the dataset under consideration.

To explore the potential of these methods to uncover previously unidentified pathway activities, we set up to perform a systematic comparison of four latent factor models recently proposed: scCoGAPS^21^, LDA^29^, scHPF^22^ and scVI^23^. We test these models on two scRNA-seq datasets from autoimmune diseases patients. The first dataset consists of single cells isolated from synovial biopsies of rheumatoid arthritis (RA) patients and further sorted into four main cell subsets (monocytes, B cells, T cells and fibroblasts, referred to as the RA dataset)^9^. The second dataset consists of single cells isolated from the kidney of systemic lupus erythematosus (SLE) patients with lupus nephritis (LN) and enriched for the leukocyte component (referred to as the SLE dataset)^7^. We evaluate the stability over iterations of the four methods across the dimensionality of the latent space by using three different metrics and highlight the predictive power of these methods to discriminate cells isolated from patients or controls. Furthermore, we assess the methods’ ability to recover gene signatures by evaluating the coverage across 13 different gene set collections. Reasoning that latent factors can be used as surrogates of pathway activities, we devise a simple method to assign gene signatures to cell clusters, thus enabling the identification of cell subsets from a functional perspective. We then extend this analytical framework to integrate ligand – receptor interactions across cell subsets. Finally, we explore the reported gene signatures and discover two previously unidentified pathways in the RA dataset: the Oncostatin M receptor signalling pathway in a subpopulation of fibroblasts and the MERTK receptor signalling pathway in a monocyte subset. We show that these signatures are potentially disease-associated, thus highlighting the power of latent factor modelling to inform the discovery of novel pathogenic pathways.

## Results

### Evaluation of latent factor models show differences in performance across tasks and latent dimensions

It has been shown that factorization solutions are not strictly convex, thus showing instability properties at different iterations^19,30^. A common heuristics to select an appropriate latent dimension is the algorithm stability across iterations^20,31^. For each method, we performed 10 iterations across 13 dimensionalities of the latent space (from k = 16 to k = 40, with step 2) and computed three stability metrics: Amari distance, silhouette score on the k-medoids-defined clusters, and the singular value canonical correlation analysis (SVCCA) score (see STAR Methods for details). We performed this evaluation for both the RA and the SLE datasets (Figure 1A and 1B). scCoGAPS and LDA emerge as the most stable methods, across latent dimensions as well as across the three chosen metrics. In contrast, scVI shows poor stability, showing better performance than scHPF only for the SVCCA score. Overall, all methods report a lower performance as the number of latent dimension increases, consistent with the increase model complexity. Running times for the four methods were found to vary considerably, with scVI and scHPF considerably faster than LDA and scCoGAPS, particularly at high dimensionality of the latent space (Supplementary figure 1A). Losses for the four methods at different values of k were also investigated and reported (Supplementary figure 1B and 1C).

**Figure 1:**
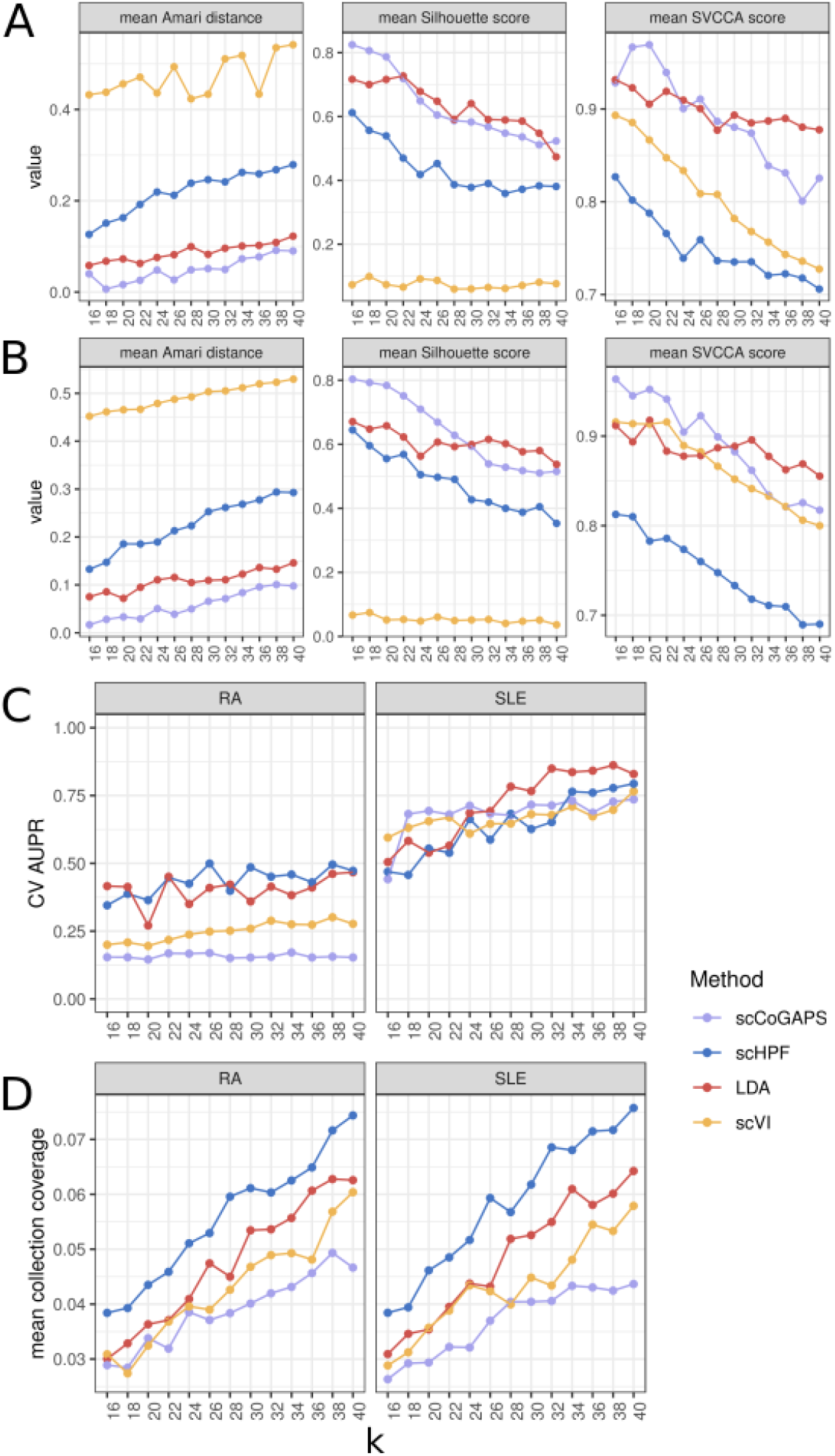
Evaluation of methods across latent dimensions. Stability metrics in A) the RA dataset and B) the SLE dataset. Y-axis reports the mean value of the metric across 10 iterations. X-axis reports k, the number of latent dimensions. C) Mean gene set collection coverage across latent dimensions, in the RA dataset and the SLE dataset. Y-axis reports mean collection coverage value, averaged across 13 gene set collections. X-axis reports K, the number of latent dimensions. Cross-validation AUPR curve in a disease – control classification task using latent variables as predictors, in the RA dataset and the SLE dataset. Y-axis reports cross-validation AUPR value across 10 iterations. X-axis reports the number of latent dimensions.

To assess whether these latent factors were useful in a classification setting to distinguish disease from control cells, we performed elastic net logistic regression with 10 fold cross-validation across dimensionalities for each of the four methods, using the latent factors as predictors and the disease state as the response variable (Figure 1C, Supplementary figure 1D and 1E). Interestingly, the performance varies considerably between datasets, but not methods. In the RA dataset, almost all methods fail to reach an AUPR > 0.5 regardless of the dimensionality of the latent space. In contrast, for the SLE dataset, all methods see a consistent increase in predictive power as the dimensionality increases, with scCoGAPS and LDA showing the best performance at low (16-24) and high (26-40) dimensionality of the latent space, respectively. Taken together, these results suggest that the dimensionality of the latent space is critical for extracting biological features related to the disease state of the cell.

The ability of latent factor models to recover biological signal is a key feature in their application to discover cellular phenotypes. Gene set enrichment analysis is a widely used approach for this task, as it allows mapping each latent variable to a specific pathway or biological process. To evaluate the methods’ ability to derive biologically meaningful gene signatures in a systematic manner, we used an enrichment approach based on heterogeneous networks^28^. Briefly, at each dimensionality of the latent space, we compute the gene set coverage score (number of unique gene sets significantly associated with each latent variable divided by the total number of gene set in the collection) for the gene set collection of interest (see Methods for details). We considered thirteen gene set collections, covering most of the known pathways and biological processes, as well as several other gene signatures (Figure 1D). As expected, for all methods we could observe an increase in the gene set coverage as the dimensionality of the latent space increases. By comparing the gene set coverage on the latent variables with the standard enrichment on clusters’ marker genes (Supplementary figure 1F), we showed that the number of significant gene sets is an order of magnitude higher for the factorization methods, pointing to a higher sensitivity in the discovery of pathway activities. Interestingly, scHPF clearly outperformed the other methods in the majority of the gene set collections in both datasets (Supplementary figure 1G and 1H). This suggests that scHPF can decompose the expression matrix in a latent space that retains the highest degree of biological signal, which prompted us to use this method for all downstream analyses.

### Systematic assignment of latent variables to cell clusters allows identification of cell types based on their phenotype or function

An open challenge in single-cell transcriptomics is the phenotypic identification of cell populations after clustering. Usually, this is performed by means of a combination of prior knowledge of cell-specific markers and gene set enrichment analysis performed on the marker genes list for each cell subset^10^. However, as latent variables provide a surrogate of pathway activities across cells, we devised a simple framework to assign each pathway to cell clusters (Figure 2A). This approach directly allows the identification of cell subsets in a function- or phenotype-driven way, avoiding the need for clustering based on marker genes. Briefly, for each gene set, we collapse redundant assignments to multiple latent variables in unique pathway activities, by means of an iterative clustering approach. Then, we regress pathway activities weights using the cluster annotation as predictors. Thus, the coefficient of each cluster represents an indicator of how important that cell subset is to explain the pathway activity. The heat map in Figure 2B is an example of the result, showing the values of the clusters coefficients for all the top significant gene sets mapped to the latent variables (using the KEGG gene set collection for the RA dataset). Interestingly, broadly defined cell populations cluster together, showing that consistent activities across different biological processes recapitulate cell lineages. Moreover, such an approach allows to discover activities that are unique to specific cell types or that are shared across different cell subsets in an unsupervised way. For instance, “B cell signalling” and “NK cell cytotoxicity” pathways (Figure 2C and 2D), show a distinct activity in the expected cell populations (see Supplementary figure 1I and 1J for an overview of the identified clusters). This strategy can also be used to discover known yet undefined cell types, such as plasmacytoid dendritic cells (Figure 2E). Finally, in the SLE dataset, we could identify a type I Interferon signature specifically active in a distinct subset of B cells and T cells, as previously reported^7^ (Figure 2F). Overall, we show that pathway activities are a powerful approach to assign cell identity to clusters based on their function or phenotype. This framework is well-suited to discover phenotypic activities that are shared across different cell types, as well as activities that are cell-type specific in both the RA and SLE datasets (Supplementary figure 2, and Supplementary tables 1 and 2, for RA and SLE respectively).

**Figure 2:**
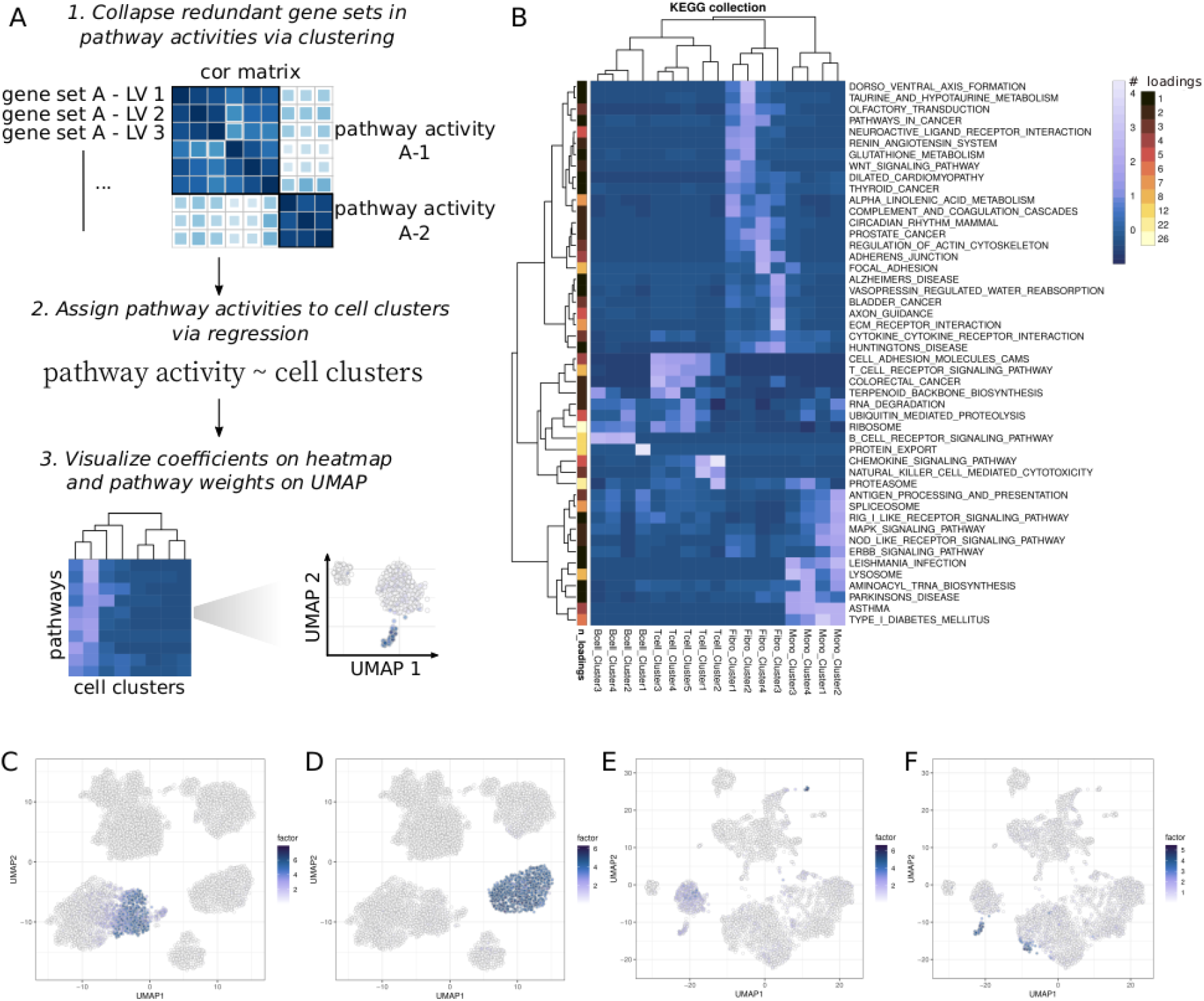
Overview of cell type-specific and activity-specific gene sets assigned to cell subsets. A) schematic of the framework to derive pathway activities and assign them to cell subsets. B) heatmap of the KEGG collection’s gene sets assigned to cell clusters in RA. The reported value represents the coefficient of the regression model and the “# loadings” colour scale represents the number of latent variables that were found significant for that specific gene set. C) factor weights for NK-cell cytotoxicity from the KEGG collection, D) factor weights for the B-cell receptor signalling pathway gene set from the KEGG collection, E) factor weights for the plasmacytoid dendritic cell signature from the C7 Immunological signature collection, F) factor weights for the type I interferon signaling from the REACTOME collection.

### Oncostatin M receptor signalling is active in specific subsets of rheumatoid arthritis fibroblasts that share a similar inflammatory profile to stromal cells from inflammatory bowel disease patients

By using latent variables as surrogates of pathway activities, we sought to discover novel pathways potentially involved in RA. We focused on Oncostatin M receptor (OSMR) signalling, whose expression level was found to be low yet widespread across fibroblast subsets (Figure 3A). However, 17 latent factors were found to be enriched for OSMR signalling-related gene sets. We collapsed these redundant gene sets in 4 pathway activities (Figure 3B) showing a distinct distribution and composition of fibroblast subsets (Supplementary figure 3A and 3B). OSMR has been recently discovered to be a driver of increased inflammatory state of stromal cells in inflammatory bowel disease (IBD)^32^. To investigate whether the fibroblast populations with high OSMR pathway activity also exhibited a similar inflammatory phenotype, we retrieved the gene signature associated with OSMR-high expression^32^ and visualized the mean gene expression in the OSMR-signaling pathway activity (Figure 3C). Interestingly, two of the OSMR activity clusters reported a high expression for the previously identified gene signature, as compared to the other two pathway activity clusters. Furthermore, these activity clusters seem to be mostly constituted by cells belonging to fibroblast clusters 1 and 2, which exhibit sub-lining markers (Supplementary figure 3C and 3D). These results suggest that the OSMR signalling pathway is active in a specific subset of sub-lining fibroblasts characterized by a gene signature that recapitulates an inflammatory phenotype observed in stromal cells associated with IBD, thus pointing to OSMR as a potential driver of such inflammatory phenotype in RA.

**Figure 3:**
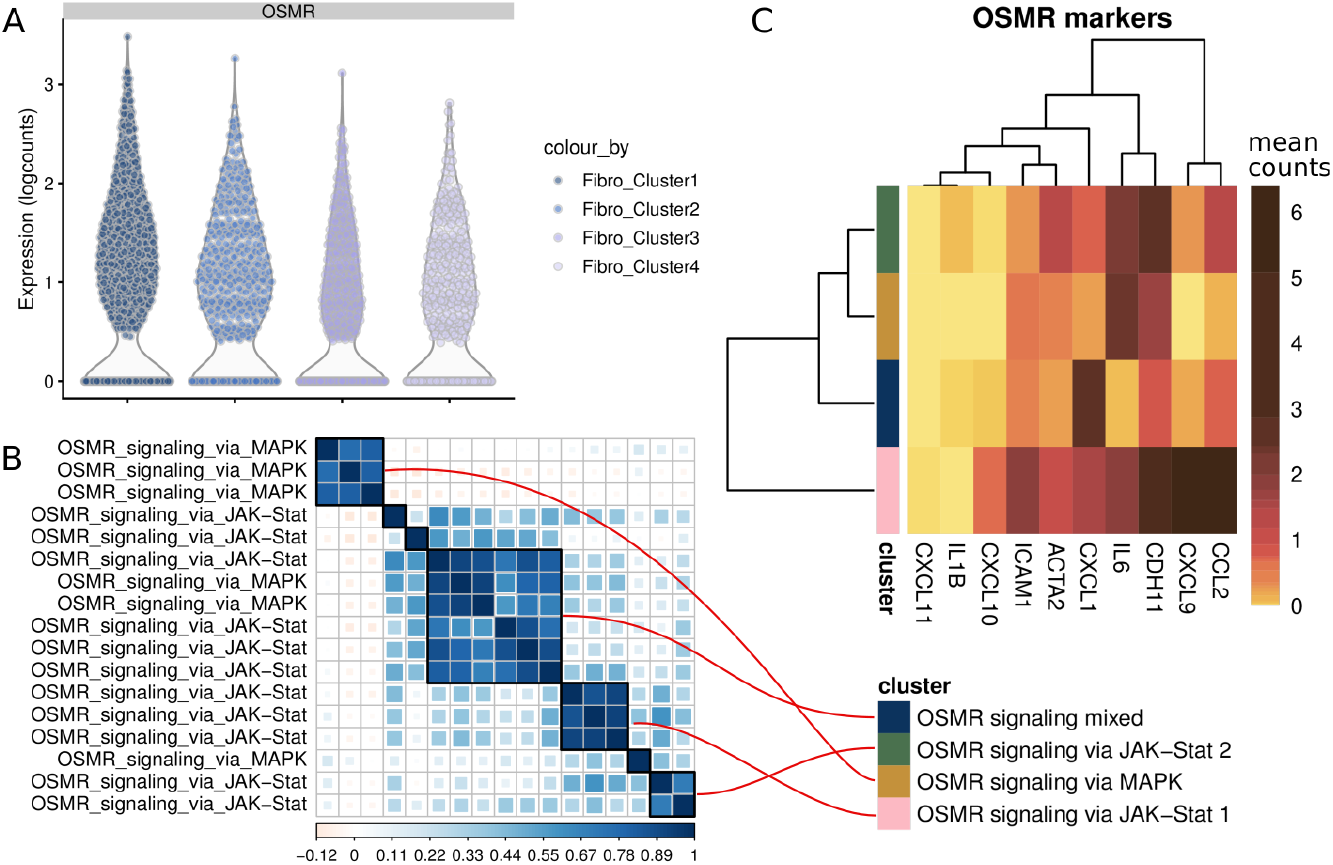
OSMR signalling pathway is specifically active in a subset of fibroblasts in RA. A) Expression levels of OSMR across fibroblast clusters, B) Correlation matrix of latent variables that maps to OSMR signalling pathways, as annotated by the METABASE gene set collection. Black frames enclose the correlation clusters that were selected to be representative of specific pathways activity, C) Mean expression level of genes found to be associated with OSMR-high-stromal cells in IBD.

### Integration of ligand – receptor interactions reveals MERTK-driven apoptotic cell clearance by a monocyte subset in rheumatoid arthritis

To further explore the potential of pathway activities to uncover novel gene signatures, we sought to integrate this information with ligand – receptor interaction analysis (see STAR Methods). In short, the expression level of proteins annotated in any interacting pair was correlated with latent variables that reported a significant enrichment for pathways where the protein is present (Figure 4A). Among the filtered cellular interactions, we found a GAS6 – MERTK link: both monocyte clusters 1 and 3 were reported to interact with GAS6 in fibroblasts and B cells subsets via MERTK (Supplementary figure 4A and 4B). Furthermore, MERTK reported a distinct expression across monocyte subsets (Figure 4B). We found MERTK-associated pathways to cluster in two main groups (Figure 4C): one with gene sets related to cell motility and cell signalling, the other one related to endocytosis and phagocytosis. As MERTK is a known marker for endocytic and phagocytic activity (particularly in the context of apoptotic cell clearance^33^), we set up to assess whether cells that were showing an endocytic-related activity reported an efferocytosis signature^34,35^. We observed that cells characterized by endocytic activity indeed recapitulated this gene signature to a higher degree as compared to the other activity clusters (Figure 4D). This cell subset, which is mostly constituted of monocytes from cluster 3 (Supplementary figure 4C, see Supplementary figure 4D for signature across monocyte clusters), was found to be depleted in RA when compared to controls (Figure 5E), suggesting that reduced apoptotic cell clearance by MERTK-signaling monocytes could be pathogenic in RA.

**Figure 4:**
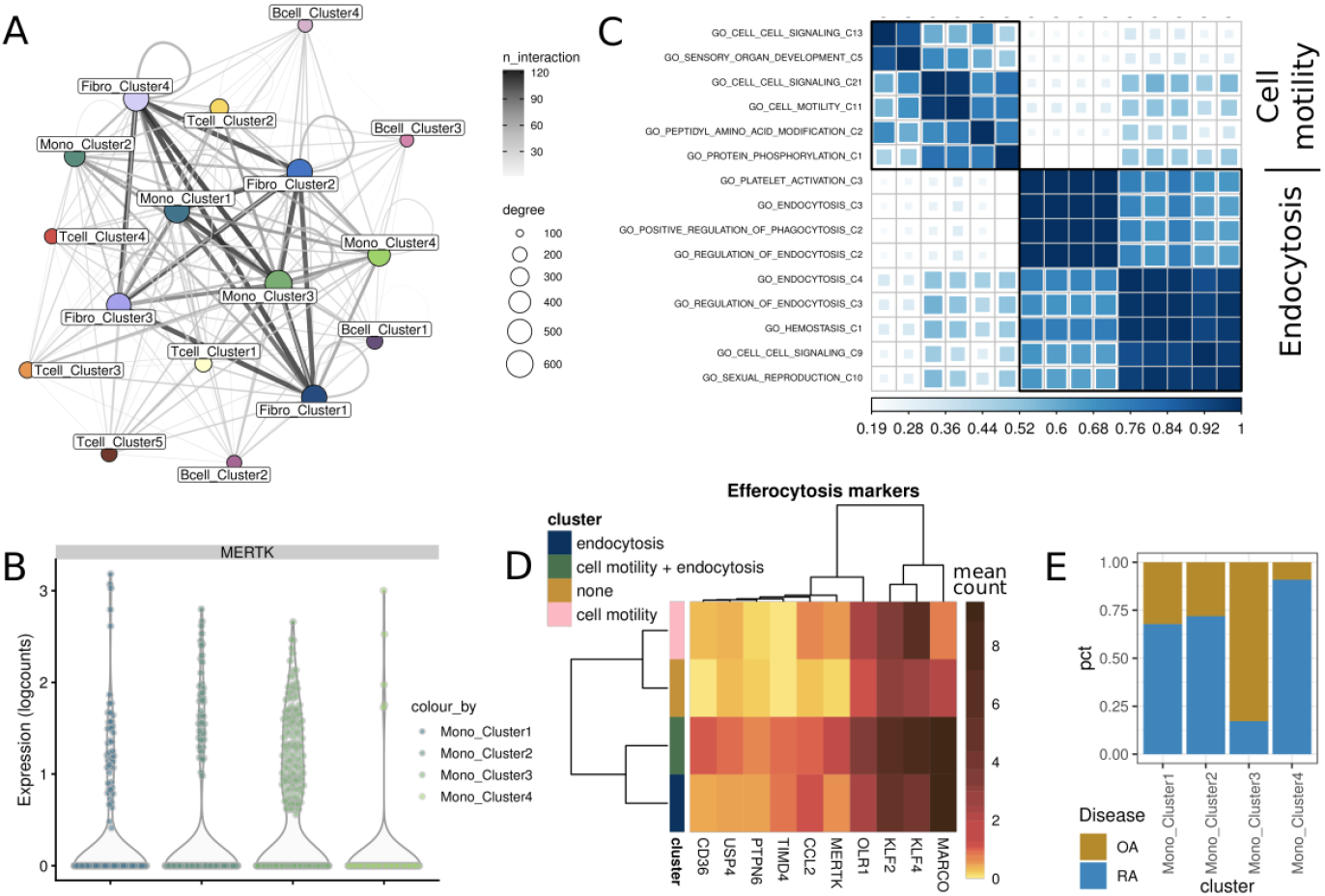
A) Ligand – receptor interaction network as computed by Cellphone Db and filtered as described in the main text. B) Expression levels of MERTK across monocyte clusters. C) Correlation matrix of latent factors that are associated with MERTK expression. Black frames enclose the correlation clusters that were selected to be representative of specific pathways activities. D) Mean expression levels of genes found to be associated with infiltrating macrophages showing an efferocytic activity. E) Proportion of cell types from disease (RA) or control (OA) across monocytes clusters.

## Discussion

Latent factor models are a flexible approach to uncover transcriptional programs in an unsupervised fashion, since they do not require prior information on the dataset structure, in contrast to differential expression and trajectory analysis. However, the strengths and weaknesses of these algorithms across different tasks are not always obvious. In addition, the biological interpretation of latent factors derived from these methods is often difficult. Here, we conducted a comprehensive evaluation of four state-of-the-art latent factor model, assessing stability, predictive power and gene set coverage of latent variables across the dimensionality of the latent space. We performed the evaluation on two autoimmune disease datasets, on which latent factor models had not been applied before. We devised a novel approach to map latent factors to pathway activities and assign these to cell subsets, showing that this framework is useful to discover previously unidentified cellular phenotypes. We focus on the activity of two signalling pathways, OSMR and MERTK, and show that their related pathway activities are distinctively triggered in specific subsets of fibroblasts and monocytes, respectively. Using literature-curated gene expression signatures, we validate the hypothesized cellular phenotypes, thus highlighting these as novel pathways involved in the pathogenesis of RA.

Benchmarking stability across iterations showed that scCoGAPS and LDA were the most robust methods, while scVI and scHPF were less stable, and showed contradictory results depending on the metric. Furthermore, in a classification setting, we showed that the predictive power of latent factors varies greatly both at different dimensionality of the latent space and at different sample sizes. The ability to recover a meaningful biological signal is an important feature of latent factor models. Here, by performing a comparison across 13 different gene set collection, we report the proportion of gene signatures retrieved by each model. As expected, the gene set coverage increases with the dimensionality of the latent space. scHPF clearly outperformed the other methods in this task and led us to select it for downstream analysis, where the aim is to discover potentially novel pathways associated with disease. Overall, the comprehensive evaluation we performed highlighted strength and weaknesses of each method in different tasks and can be used as a starting point for method selection depending on the user needs (stability, predictive power, discovery of biological pathways).

In order to leverage the latent factors as surrogates of pathway activities and aid the identification of cell types, we devised a novel framework to collapse redundant gene sets into pathway activities and assign these to cell clusters. We show that such an approach is able to retrieve known gene signatures and previously identified cellular phenotypes (such as the type I interferon signalling active in specific subsets of B cells and T cells in SLE^7^). As shown with the plasmacytoid dendritic cell subset in SLE, we also show that it can be used to identify rare cell subpopulations based on known cell identity signatures without using pre-specified marker genes. As it relies heavily on gene sets annotation, the quality of the resulting pathway activities is inevitably tied to the quality of the original gene sets in the collection. As such, some of the identified pathway activities might be false positives. One example of this is the olfactory transduction gene set from the KEGG collection, which was found to be significantly associated with 64 latent factors, further collapsed in 19 pathway activities. Therefore, we suggest that hypothesis-driven explorations of the pathways activities assignments are needed to draw meaningful interpretations, coupled with orthogonal analysis to identify cell types. While we focused on two autoimmune disease cohorts, the described framework is generally applicable to any scRNA-seq datasets and provides an intuitive way to directly define cell subpopulations based on their function or phenotype, without having to rely on cell marker genes and iterative clustering procedures.

Among the pathways activities that were not previously reported by the authors of the original publication^9^, we noticed the Oncostatin M signalling pathway to be frequently associated with different fibroblast clusters. Interestingly, Oncostatin M has been recently reported to be a key driver of intestinal inflammation in IBD and to be associated with response to anti-TNF alpha therapy^32^. We showed that fibroblast subsets that show an OSMR signalling pathway activity also express a high level of a gene signature found to be associated with OSMR high expression in stromal cells from the intestine of IBD patients. This points to the potential involvement of Oncostatin M in establishing the inflammatory microenvironment in RA patients and suggests the pathway might be similarly dysregulated in RA and IBD. Interestingly, cells that report such specific OSMR signaling activity mainly belong to sublining fibroblast subsets. Since the same fibroblast subpopulation has also been recently associated with a specific inflammatory phenotype^36^ (as opposed to a more cartilage degradation phenotype of the lining subsets) this points to the potential involvement of OSMR in driving this specific inflammatory profile in both IBD and RA.

To further explore the potential of pathways activities to uncover novel cellular phenotypes in RA, we integrated this approach with ligand – receptor interaction analysis and identified a GAS6 – MERTK link. Interestingly, we found that MERTK expression correlates with two main groups of pathway activities: one which entails gene sets related to cell motility and cell signalling, and one which entails gene sets related to endocytosis and phagocytosis. To support the hypothesis that monocytes that display the latter might be involved in apoptotic cell clearance, we evaluated the expression of genes related to an efferocytosis signature^34,35^. Interestingly, we were able to show that this monocyte subset well recapitulates such a signature. The subset mostly corresponds to a monocyte cluster which is reported to interact via MERTK with a number of infiltrating lymphocytes and fibroblasts, and is depleted in disease. These results replicate the clusters reported by the authors (SC-M2 and SC-M3)^9^, who were, however, not able to assign a function to these monocyte subsets. Interestingly, a recent paper^37^ seems to support the presence of an infiltrating macrophage subpopulation that exhibits such a resolving phenotype, substantiating our findings. Overall, these results highlight how integrating cellular interaction analysis and pathway activities might help in the identification of previously unidentified pathways dysregulated in specific cell subsets.

In conclusion, our benchmarking results highlight strengths and weaknesses of latent factor models applied to scRNA-seq data for different tasks. We focused our work on the application of these methods for the purpose of signature discovery, yet latent factor models have been implemented for a variety of purposes (such as denoising^38,39^ and multi-omics integration^40,41^). We propose a framework that makes use of the full spectrum of latent variables and allows to directly define cell subtypes based on their function or cellular phenotype. Finally, we apply the framework to scRNA-seq data from synovial biopsies of RA patients and implicate two novel pathways in the disease pathogenesis: activation of OSMR signalling in a subset of inflammatory fibroblasts and dysregulation of the GAS6 – MERTK axis in an efferocytic monocyte subtype.

## STAR Methods

### Datasets

The two datasets that we considered for the analysis were collected and provided by the NIH Accelerating Medicine Partnership (AMP) consortium, via Immport. The RA dataset consisted of cells from 3 osteoarthritis (OA) patients and 22 rheumatoid arthritis (RA) patients, isolated from the synovium and further sorted via fluorescent activated cell sorting (FACS) into 4 subpopulations: fibroblasts, monocytes,B cells and T cells. The SLE dataset consisted of cells from 3 healthy donors and 15 lupus nephritis (LN) patients, a complication of SLE which involves the kidney. Cells were isolated from the kidney and further sorted for CD45+ markers, in order to retain an enriched population of leukocytes.

### QC, feature selection and clustering

scRNA-seq raw counts were downloaded from ImmPort (study number SDY998 and SDY999 https://www.immport.org/shared/home). QC was performed, in both datasets, with the *Scater* package^42^ with the following thresholds: cells had to have > 1000 and < 5000 UMI counts to filter out potential doublets as well as dead cells. Also, the percentage of mitochondrial reads had to be below 0.25. Genes had to have at least 3 UMI in at least 3 cells. Further gene filtering was performed by means of the *deviance*^43^ definition. 6000 genes and 3000 genes were selected for the RA and SLE dataset respectively. Clustering was performed with SNN-cliq^44^ via the *scran* package^45^ and the selection of the clustering granularity was aided by the *clustree* package^46^. UMAPs^47^ of the 2 datasets and respective clusters are reported in Supplementary figure 1I and 1J.

### Latent factor modelling algorithms

Four methods were selected for evaluation: scCoGAPS^21^, LDA^29^, scHPF^22^ and scVI^23^. The number of dimensions for the latent space (16 to 14 latent variables, step 2) has been selected for efficiency and for consistent evaluation across models (i.e. a running time suitable to all models, see Supplementary figure 1A). Here, the loading matrix refers to the matrix latent variables v. genes, whereas the factor matrix refers to the matrix cell v. latent variables. At each dimensionality of the latent space, we run the methods for 10 iterations. For scCoGAPS, we employed a parallelization approach as suggested by the authors^48^, and set the maximum iteration parameter to 7000. For scVI we made use of the implementation of the linear decoder^49^, so that the decoder weight matrix could be used as a surrogate of the loadings matrix. Model hyperparameters were set based on those selected for datasets of similar size and characteristics, as reported by the authors^23^. For LDA we employed hyperparameters as described in a similar use case^27^. For scHPF, we employed hyperparameters as suggested by the authors^22^.

### Stability evaluation

Stability evaluation was performed across iterations and latent dimensions. Three metrics were used:

- *Amari-like distance*^30^: a correlation-based metrics, that we computed for each pairs of iterations. The mean of all the comparisons is reported.
- *Silhouette score:* similar to Kotliar et al.^20^, the silhouette score was calculated based on the clusters of the concatenated loading matrices. Briefly, ten loading matrices, computed for each iteration, were concatenated and further clustered with a k-medoids approach. The number of clusters was set equal to the number of latent dimensions. The mean of all the comparisons is reported.
- *SVCCA*^50^: similar to Way et al.^28^, SVCCA computes singular value decomposition on two loading matrices, and then perform canonical correlation analysis, to align matching components and derive correlation coefficients between them. The mean of all the comparisons is reported.

### Gene set coverage evaluation

Gene set coverage score was computed by means of heterogeneous networks^51^, as described in Way et al.^28^. Briefly, we made use of heterogeneous network made available by Way et al. or generated as part of this study from several gene set collections (MSigDB^52^, KEGG, REACTOME, Biocarta, MetaBase^18^(Clarivate Analytics MetaBase^®^ version 6.15.62452), WikiPathways^53^). We also generated respective shuffled networks in order to calculate a Z-score for each gene set – latent variable pair. We then converted the Z-score to a p-value and filtered using a Bonferroni correction (p-value threshold defined as 0.01 divided by the number of latent variables for the specific model – latent space dimensionality pair). For each gene set collection and each latent variable, the top gene set was selected to be mapped to that latent variable. Ultimately, for each model, we would have an equal number of gene sets according to the dimensionality of the latent space. The number of unique gene sets was then used to calculate the coverage score (number of unique gene sets divided by the number of total gene sets in that collection). In Supplementary figure 1G and 1H, the mean collection coverage value was calculated across iterations for each dimensionality of the latent space, for the RA and SLE datasets, respectively. In Figure 1C, the mean collection coverage, across collections, was calculated for the best iteration at a fixed k, for each method. The selection of the best (i.e.: run with lowest reported loss) factorization result for a given k, was used for all downstream analyses. To perform the comparison with gene lists generated by means of differential expression across clusters, we assumed that each cluster-specific signatures represent a latent feature. We used −log10 (p-value) as gene weights, so to have a genes by clusters table, which we then used as input for the gene set coverage evaluation. Since the number of clusters is 17, we showed a comparison of gene set coverage with the other methods at k=16 and k=18 (Supplementary figure 1F).

### Gene set assignment to cell clusters

After matching each latent variable to its most significant gene set in each collection, we reasoned that we could use the latent variable as a surrogate of the respective pathway activity. Furthermore, we decided to use all the latent variables derived by the models, without selecting a single dimensionality of the latent space. For this and downstream analysis, we made use of the latent variables as computed by *scHPF*, selecting the best run (as assessed by the loss) for each dimensionality of the latent space. To account for the duplicated instances of the gene sets (same gene set mapped to multiple latent variables), we implemented a simple iterative algorithm. If a gene sets mapped to 3 or more latent variables, we would cluster the correlation matrix of the latent variables and compute the mean Silhouette width at different cut (H) of the hierarchical tree (where 1<H<= # latent variables – 1). Then, for all the clusters that had a mean Silhouette width > 0.5 we collapsed them by computing the medians of the weights of the latent variables belonging to the respective clusters. If the Silhouette width <= 0.5 or the gene set mapped to only 1 or 2 latent variables, we just collapsed all the latent variables mapping to that gene set into the mean. This would provide us with collapsed latent variables for downstream analysis that we refer to as pathway activities. If the pathway activities were split during this clustering step, then it would be reported with a unique identifier (e.g. pathway_A_C2, representing the additional instance of pathway_A).

Furthermore, we devised an approach to assign these pathway activities to cell clusters: each pathway activity was set as the response variable in a regression setting where the cluster labels function as the predictors. Thus, a cell cluster that comprises several cells that have a high weight for that specific latent variable, would also be assigned a large coefficient. The pathway activities were standardized before the regression step. The pathway activities reported in Figure 2B represent the cell clusters’ coefficients for all the pathway activities (see Supplementary figure 2 for the remaining collections). If the gene set was collapsed in several different pathway activities, those assignments are also reported.

### Gene signature analysis

#### Oncostatin M receptor signature

Pathways mapping to OSMR signalling were clustered and processed as described above. Median weights of the four selected clusters (Figure 3B) were used to filter cells (weights > 0.5 quantile). The mean gene expression signatures for the selected genes were calculated for cells that were specifically filtered in one of the four clusters but did not intersect with others. The expression of the same gene list was also reported for the fibroblast clusters (Supplementary figure 3C). The gene signature was retrieved from West et al.^32^.

### MERTK signature

CellPhoneDB^17^ was run with following parameters: *statistical_analysis, --iterations 5000, --threshold 0.2*. All gene sets activities (i.e.: weights of the latent variables) where MERTK is present were correlated with MERTK expression levels, in monocyte cells. All gene sets that showed an R^2^>0.3 were then filtered and the correlation matrix was clustered (Figure 4B). Two clusters were selected: the cell motility cluster and the endocytosis cluster. Medians of the latent factor weights were calculated for these two clusters, and cells were filtered for their respective activity (as described above). Cells that showed the activity for either, both or none of the two clusters were filtered and mean expression values were reported for genes retrieved from a gene from Waterborg et al. 2018^34^ and Robert et al. 2017^35^. Supplementary figure 4D shows the mean expression of genes belonging to the efferocytosis signature based on clustering. Supplementary figure 4E shows the composition of the different pathway activities in terms of monocyte clusters.

### Visualization and software

All visualizations were produced with the ggplot2 package^54^. Methods and comparison analysis were wrapped in a Snakemake pipeline^55^

### Code availability

All code of the analysis can be found at https://github.com/giovp/latent_factors_autoimmune

## Abbreviations

AMP: Accelerating Medicine Partnership
FACS: fluorescent activated cell sorting
IBD: inflammatory bowel disease
ICA: independent component analysis
LN: lupus nephritis
NMF: negative matrix factorization
OA: osteoarthritis
PCA: principal component analysis
RA: rheumatoid arthritis
SLE: systemic lupus erythematosus
SVCCA: singular value canonical correlation analysis

## Acknowledgements

We would like to thank Jonas Zierer, Christine Huppertz, Grigory Ryzhakov, Dominik Hartl, Stephan Spiegel, James Rush and Richard Siegel for helpful discussions and comments on the manuscript and Patrick Dunn for helping with data access.

